# Explainable Machine Learning for Preoperative Relapse Prediction in Molecularly Stratified Endometrial Cancer: A Single-Center Finnish Cohort Study

**DOI:** 10.1101/2025.11.06.686680

**Authors:** Sergio Vela Moreno, Masuma Khatun, Annukka Pasanen, Ralf Bützow, Andres Salumets, Mikko Loukovaara, Vijayachitra Modhukur

## Abstract

Relapse risk in endometrial carcinoma (EC) is strongly influenced by molecular subtype, yet current WHO/ESGO classifications rely on postoperative data, limiting their utility for preoperative decision-making. We developed and compared interpretable machine learning (ML) models to predict relapse timing (none, ≤6 months, >6 months) using exclusively preoperative multimodal data. In a retrospective cohort of 784 EC patients, we integrated clinicopathological, molecular, immunohistochemical, and systemic biomarkers and constructed four feature strategies: (1) Traditional (clinicopathology), (2) ESGO (guideline risk groups), (3) TP53 + MMRd (high-risk biology), and (4) POLE (low-risk biology). Classifiers (Random Forest (RF), Support Vector Machine (SVM), k-Nearest Neighbors (KNN), Gradient Boosting (GBM)) were trained with leakage-safe preprocessing and in-fold resampling; performance was evaluated via area under the curve (AUC), accuracy, recall, and F1 score, and interpretability via SHapley Additive exPlanations (SHAP). The RF-based Traditional model achieved the highest overall performance (F1 = 0.895, AUC = 0.84), while the GBM-based POLE model showed superior sensitivity (F1 = 0.886, AUC = 0.842). SHAP identified ARID1A loss, elevated CA125, thrombocytosis, and p16 expression among key predictors of relapse; while overlapping high-risk features across models included advanced stage, deeper myometrial invasion, elevated CA125, and positive cytology. These biologically coherent, explainable predictions support individualized risk stratification and may enhance preoperative decision-making, particularly for aggressive histology and high-risk molecular subtypes.

**Graphical abstract:** 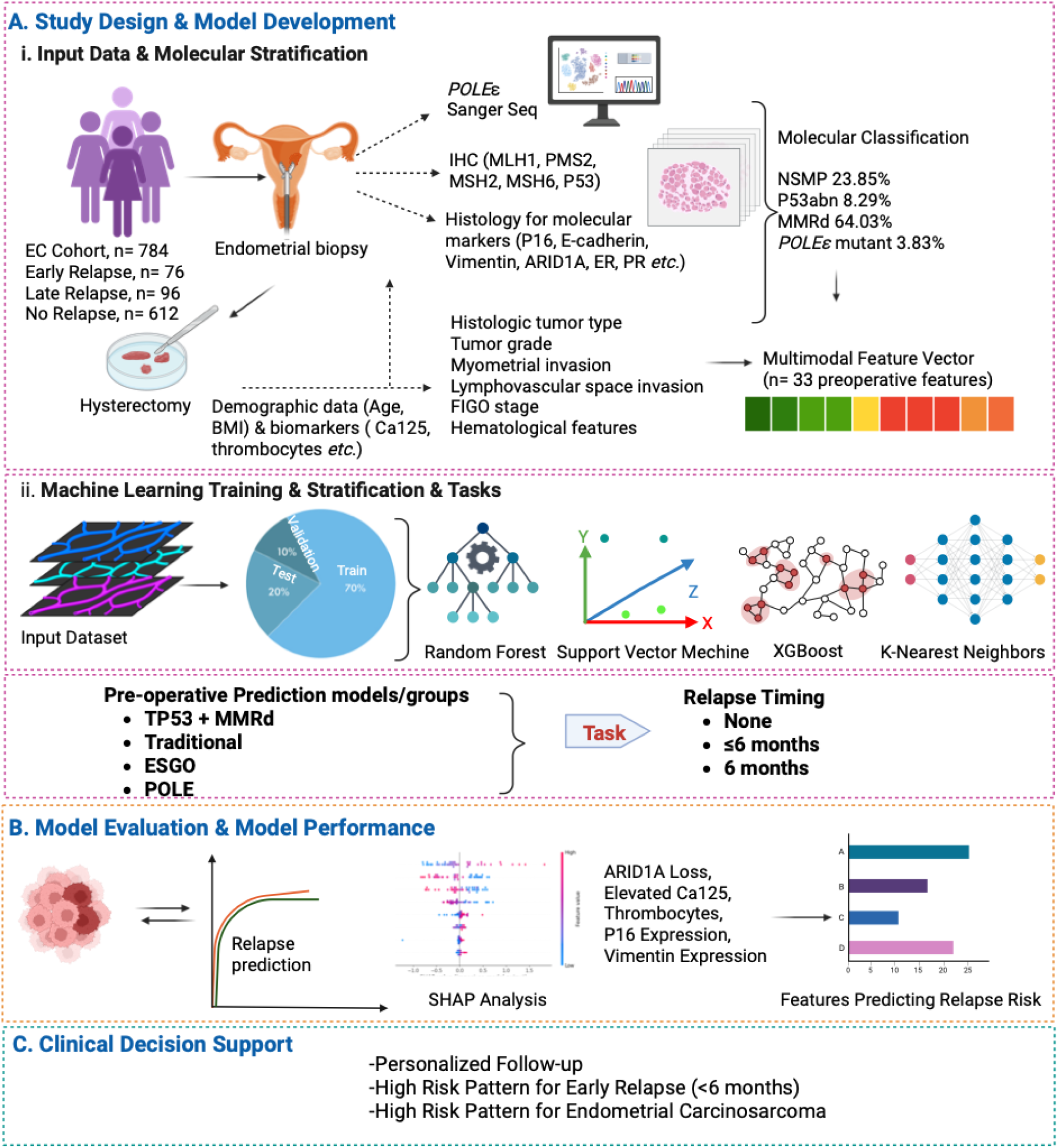

**Workflow for Machine Learning (ML)–based relapse prediction in endometrial cancer.** The schematic figure outlines the study pipeline from patient inclusion to clinical application. (A) A retrospective cohort of 784 EC patients was analyzed, integrating clinical, demographic, biomarker, and molecular data into a multimodal feature set. Patients were stratified into four molecular subgroups: NSMP, p53abn, MMRd, and *POLE*mut. Multiple ML algorithms (Random Forest, SVM, XGBoost, k-NN) were trained to predict relapse timing. (B) Model performance was evaluated using area under the curve (AUC) and accuracy metrics, with SHapley Additive exPlanations (SHAP) analysis applied to identify key predictive features across models. (C) SHAP-based interpretation was used to support individualized relapse risk stratification, enabling potential clinical decision-making for surveillance and therapy.

**Highlights:**

- Pre-operative XAI models predict relapse timing in EC with an AUC of up to 0.842.
- The traditional model achieves a top accuracy of 0.797 using 22 features, while POLE maximizes sensitivity at 0.886.
- SHAP explanations identify class-specific drivers such as stage, LVSI, size, cytology, CA125, and PR.
- Early Relapse is associated with burden and aggressiveness, while Late Relapse relates to the spread and size of EC, while No Relapse indicates an inverse profile.
- Transparent outputs facilitate risk-aligned surveillance and treatment planning before surgery.

## 1. Introduction

Endometrial cancer (EC) is the most common gynaecologic malignancy in developed countries, with incidence and mortality projected to increase by 55% by 2030 (Clarke et al., 2022; Onstad et al., 2016), largely driven by obesity and metabolic syndrome (Onstad et al., 2016; Siegel et al., 2022). Despite advances in adjuvant therapies, including immunotherapy trials (RUBY, GY018, and DUO-E) (Bogani et al., 2024; Yang and Wang, 2019), 15–20% of patients experience relapse, often in the vaginal vault, pelvis, peritoneum, or distant organs, with a poor prognosis and limited treatment options (Beavis and Fader, 2022; Crosbie et al., 2022; Tuninetti et al., 2024). Endometrial carcinosarcoma, a rare but highly aggressive subtype accounting for 5-6% of ECs (Capozzi et al., 2024), exhibits relapse rates of 40–60% and high mortality (Cantrell et al., 2015).

Traditional risk stratification relies on clinicopathological features such as age, tumor grade, FIGO (International Federation of Gynecology and Obstetrics) stage, and lymphovascular space invasion (LVSI) (Concin et al., 2021; Zwahlen et al., 2016). However, these parameters often lack reproducibility and prognostic accuracy, particularly in high-risk or recurrent cases (Oaknin et al., 2022). Molecular classifiers, including The Cancer Genome Atlas (TCGA) and its clinical surrogate, The Proactive Molecular Risk Classifier for Endometrial Cancer (ProMisE), have refined EC stratification into four distinct subtypes: *POLE*-ultramutated (*POLE*mut, favorable, low relapse risk), mismatch repair-deficient (MMRd, higher relapse risk), p53-abnormal (p53abn, poor prognosis, higher relapse risk), and no specific molecular profile (NSMP, intermediate) (Lee et al., 2025; Levine, 2013). These are now integrated into the 2023 FIGO staging, which recognizes *POLEmut* tumors as favorable and p53abn as adverse, even in early-stage disease (Gaffney et al., 2024; Talhouk et al., 2017).

Despite these advances, relapse prediction remains challenging, particularly in NSMP, p53abn, and carcinosarcoma subgroups (León-Castillo et al., 2025; W Glenn McCluggage et al., 2023). Existing models often fail to incorporate comprehensive molecular and clinical variables (Urick and Bell, 2019), while symptom-based surveillance may miss early asymptomatic relapses that are linked to poorer outcomes (Backes et al., 2018; Kommoss et al., 2018). Post-relapse survival varies significantly: 43 months for MMRd, 39 for NSMP, and only 10 for p53abn, underscoring the urgent need for improved predictive tools (León-Castillo, 2023; Siegenthaler et al., 2022). Biomarkers (*e.g.*, ARID1A (Liu et al., 2017), p16 (Murali et al., 2019), β-catenin, E-cadherin (Schlosshauer et al., 2002), and systemic markers (e.g., CA125, platelet count) (Njoku et al., 2022; Schlosshauer et al., 2002) show promise but lack integration into multivariable risk models.

Machine learning (ML) continues to show promise in enhancing relapse prediction in EC, particularly when integrated with molecular classification. Ensemble models (e.g., AdaBoost, XGBoost, and Random Forest, RF) combined with interpretability tools like SHAP (SHapley Additive exPlanations) have boosted both predictive accuracy and clinical applicability. For instance, the TJHPEC model achieved an area under the curve (AUC) of 0.93 using routine clinical features across 1,935 patients (Wang et al., 2022), while radiomics-based models leveraging preoperative CT scans reached AUCs up to 0.90 (Camelia Alexandra Coada et al., 2023). Molecularly informed models like im4MEC correlated strongly with 5-year relapse-free survival (Fremond et al., 2023) and NU-CAT predicted progression and relapse with 75% accuracy (Zheng et al., 2023). Additional approaches include Random Forest (RF)-based predictors of high-grade EC (AUC 0.85) (Piedimonte et al., 2022), biomarker-integrated nomograms (Cong et al., 2023), deep learning on haematoxylin and eosin (H&E)-stained slides (Coudray et al., 2018; Fu et al., 2020), and the HECTOR model for distant relapse prediction (Volinsky-Fremond et al., 2024). However, few models have been validated in molecularly stratified cohorts, and preoperative, multi – class prediction of relapse timing, particularly in high-risk EC and carcinosarcoma, remains underexplored.

To address these gaps, we developed and compared interpretable ML models for preoperative relapse prediction of relapse timing (No Relapse, ≤6 months, >6 months) in EC, including carcinosarcoma. Four complementary approaches were implemented: 1) Traditional clinicopathological model, (2) ESGO guideline-based model, (3) Tp53 + MMRd biology-driven high-risk model, and (4) POLE low-risk feature strategies. By integrating multimodal preoperative data, including molecular classifiers, biomarkers, and clinicopathological features, we sought to evaluate trade-offs between accuracy, sensitivity, and clinical utility, while ensuring model interpretability through SHAP (SHapley Additive exPlanations).

## 2. Materials and Methods

### 2.1 Study Design and Patient Cohort

This retrospective study included 784 patients with stage I–IV EC who underwent hysterectomy at Helsinki University Hospital between 2007 and 2013. Ethical approval was obtained from the Helsinki University Hospital Institutional Review Board (HUS/491/2021) and the Finnish Medicines Agency (FIMEA/2021/005153). Informed consent was waived for this retrospective cohort.

Clinicopathological data were retrieved from institutional records. Staging followed the FIGO 2009 guidelines (Pecorelli, 2009). Tumours were classified into molecular and clinicopathological risk groups using the ESGO/ESTRO/ESP 2021 guidelines (Concin et al., 2021). Tumours were classified into low, intermediate, high-intermediate, high, and advanced-metastatic risk categories based on molecular subtype and clinicopathological factors, including histology, grade, depth of myometrial invasion, LVSI, and FIGO stage. The guidelines do not assign risk categories to stage I–IVA MMRd and NSMP clear cell carcinomas with myometrial invasion, or to stage III–IVA *POLE*mut tumours, due to limited supporting data. In this study, these tumors were classified as high-risk. LVSI was assessed using a three-tiered system: none, focal, or substantial (Concin et al., 2021). Relapse status was obtained from hospital or referral centre records, and cytology from peritoneal washings taken during surgery.

### 2.2 Preoperative Clinical and Biomarker Assessment

Preoperative data included American Society of Anaesthesiologists (ASA) physical status scores extracted from anaesthesia records and standardized to the 2014 classification system (Kolehmainen et al., 2019). Patients who were current smokers, had a BMI of 30–40 kg/m², or had well-controlled diabetes were assigned ASA II, while those with severe obesity (BMI ≥40 kg/m²) were classified as ASA III. Hematologic parameters were obtained from pre-treatment blood count using photometry, impedance, and flow cytometry. Anaemia was defined as haemoglobin (Hb) <117 g/L, leukocytosis as WBC > 8.2×10⁹/L, and thrombocytosis as platelets > 360×10⁹/L (Nordin et al., 2004). Serum CA125 levels were measured via chemiluminescent microparticle immunoassay on the Abbott Architect 2000i system, with values >35 U/mL considered elevated (Bast et al., 1981). A tumor size threshold of 25 mm was applied based on prior evidence linking it to relapse risk (Sozzi et al., 2018).

### 2.3 Molecular Classification and Immunohistochemistry

Multicore tissue microarrays (four tumor cores/case) were constructed following established protocols and scanning using the 3D Histech Pannoramic 250 Flash II. Digital images were reviewed via WebMicroscope. Immunohistochemical (IHC) scoring was performed by a pathologist blinded to clinical outcomes (A.P.), with equivocal cases confirmed by a second pathologist (R.B.) (Khatun et al., 2025; Khatun et al., 2021; Pasanen et al., 2016). Molecular classification followed WHO guidance, assigning tumors to MMRd, P53abn*, POLE*mut, or NSMP. In cases with overlapping features, classification was based on the prognostically dominant alteration (Höhn et al., 2021; León-Castillo et al., 2020; Pasanen et al., 2021; Pasanen et al., 2020). Fresh-frozen tumor samples were collected for POLE mutation analysis, with exonuclease-domain hotspot mutations confirmed by targeted sequencing (exon 9, 13, 14) (Pasanen et al., 2019). IHC panels, scoring thresholds, and assay details are summarized in **Supplementary Table S1.**

### 2.4 Explainable Machine Learning Framework

To predict relapse risk, supervised machine learning pipelines were implemented to assess relapse risk, categorized into multi-class, namely, No Relapse, ≤6 months, or >6 months. Data was pre-processed by eliminating features with >30% missing values, and only pre-operative variables were retained. Missing values were imputed using median/mode substitution with the *na.roughfix* function in the R package *randomForest*. Variables were one-hot encoded, normalized, and outliers capped using interquartile range (IQR) thresholds. After this preprocessing, the final feature sets were selected for every molecular dataset (28–29 variables depending on the molecular dataset).

To capture different perspectives of relapse prediction, four complementary models were developed. The Traditional model incorporated established clinicopathological features (FIGO stage, grade, histology, LVSI, tumor size, receptor status) and served as the benchmark reflecting current practice. The ESGO model applied the 2021 ESGO/ESTRO/ESP risk classification as a guideline-based comparator. The TP53 + MMRd model targets two molecularly defined high-risk subgroups (TP53-abnormal and MMR-deficient) with a compact, biology-driven feature set. Finally, the POLE model focused on the biologically distinct *POLE*mut subgroup, typically associated with excellent prognosis, to assess whether subgroup-specific modeling improved discrimination.

For each strategy (traditional, ESGO, TP53 + MMRd, POLE), a dedicated data frame was generated and split into 70:20:10 train-test-validation sets. Recursive feature elimination (RFE) was performed using 10-fold cross-validation (CV), optimizing for the metric F1 score in risk prediction, given the significant class imbalance between classes. Feature selection was implemented manually using the *tidymodels* R package. Tumor size thresholds were iteratively optimized to improve accuracy (Sozzi et al., 2018).

### 2.5 Model Development and Evaluation

Four supervised classification algorithms were evaluated: Random Forest (RF), Support Vector Machines (SVM), k-Nearest Neighbors (KNN), and Gradient Boosting (GBM). Models were optimized using grid search with 10-fold stratified cross-validation. Class imbalance was addressed through SMOTE, under-sampling, and oversampling, which were applied within each field.

All models for relapse risk were optimized for AUC, a threshold-invariant metric that minimizes the effect of class imbalance. Evaluation metrics included:

● Accuracy = (TP + TN) / (TP + TN + FP + FN)
● Recall/Sensitivity = TP / (TP + FN)
● F1-score = 2TP / (2TP + FP + FN)
● AUC for each class, calculated as the integral under the ROC curve (representation of True Positive Rate (TPR) (TP / (TP + FN)) and False Positive Rate (FPR) (FP / (FP + TN))).
● Precision-Recall area under the receiver operating characteristic curve (PR-AUC) for each class, calculated as the integral under the Precision-Recall curve (representation of Precision (TP / (TP + FP)) and Recall (TP / (TP + FN)).

TP=True Positive, FP=False Positive, TN=True Negative, FN=False Negative

All models were implemented in R 4.5.0 using caret 7.0-1, randomForest 4.7-1.2, dplyr 1.1.2, gbm 2.2.2, kernelshap 0.7.0, and shapviz 0.9.7. Analyses were performed on a 64-core Intel Xeon server (256 GB RAM, Ubuntu 20.04 LTS).

### 2.6 XAI-Based Model Interpretability

To enhance clinical trust and applicability, SHAP was employed to quantify feature contributions to predictions (Lundberg et al., 2017). SHAP values were computed using kernelshap with k-means background sampling (m=50, 1,000 samples per observation). Class-specific SHAP values were visualized using shapviz, enabling identification of key predictors for each relapse category (Early, Late, No Relapse). Implementations were performed using R packages *kernelshap* and *shapviz*. A summary of the ML algorithms and parameters is shown in **Table 1**.

**Table 1.**
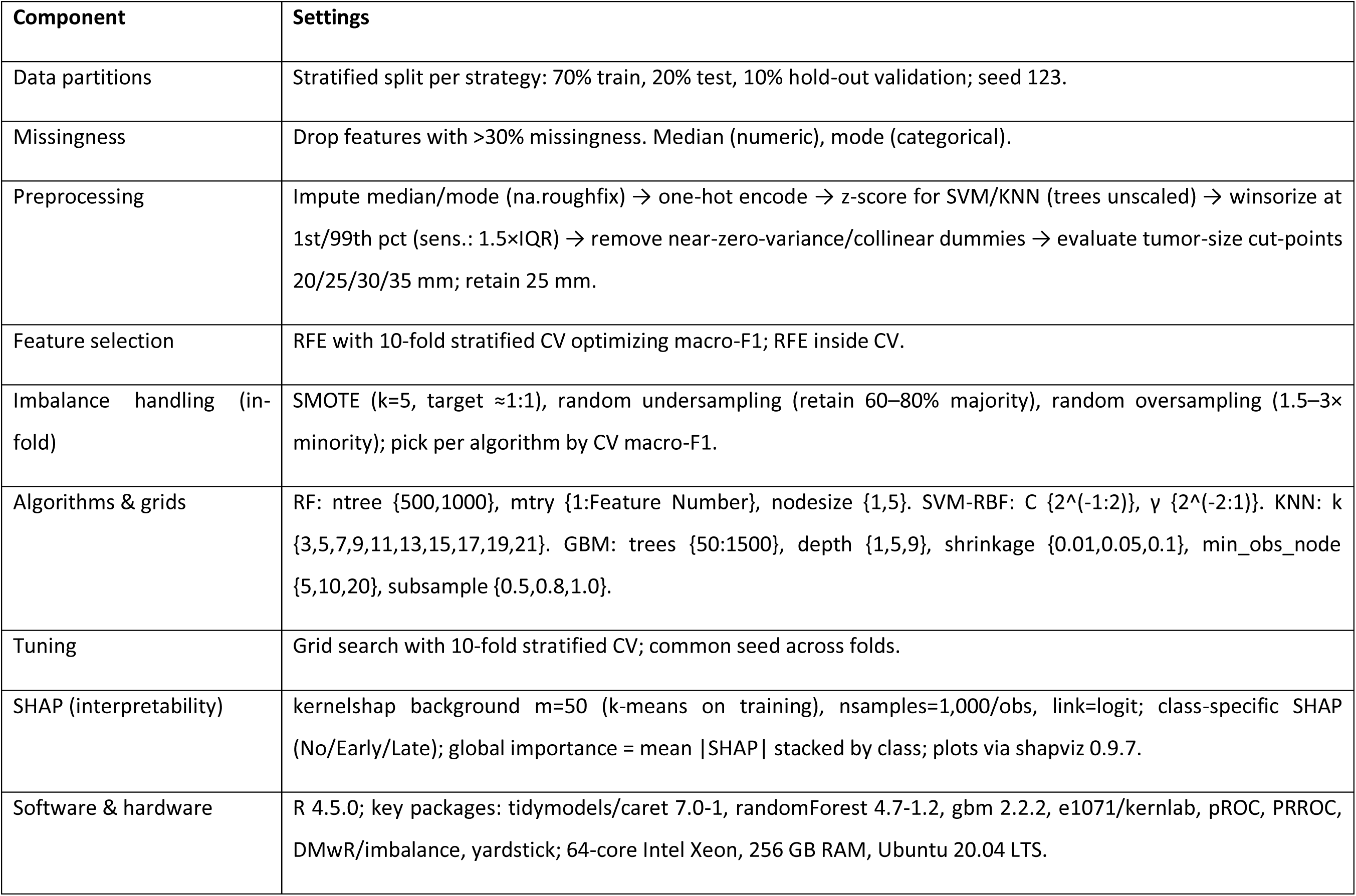
Summary of the Machine Learning algorithm.

## 3. Results

### 3.1 Patient Cohort and Molecular Stratification

A total of 784 patients with EC were included, of whom 172 (22%) experienced relapse — 76 Early (≤6 months) and 96 Late (>6 months) relapse. The remaining 612 patients (78%) remained relapse-free. Patients were stratified into four molecular subgroups: MMRd (64.0%), NSMP (23.9%), p53abn (8.3%), and POLEmut (3.8%). Thirty-three preoperative features were integrated into a multimodal feature vector for ML analyses **(Graphical abstract**).

Relapsed cases were significantly enriched in advanced FIGO stages (II–IV), non-endometrioid histology (serous and carcinosarcoma), and positive LVSI. Tumors >25 mm, positive peritoneal cytology, deep myometrial invasion, and molecular alterations (p53abn, MMRd) were more frequent in relapsed patients. On the other hand, POLEmut tumors showed the lowest relapse rates. Additional relapse-associated features included p16 positivity, E-cadherin loss, vimentin expression, ARID1A loss, elevated CA125, and increased thrombocyte and leucocyte counts. Full demographic and clinicopathological comparisons are presented in **Table 2**.

**Table 2.**
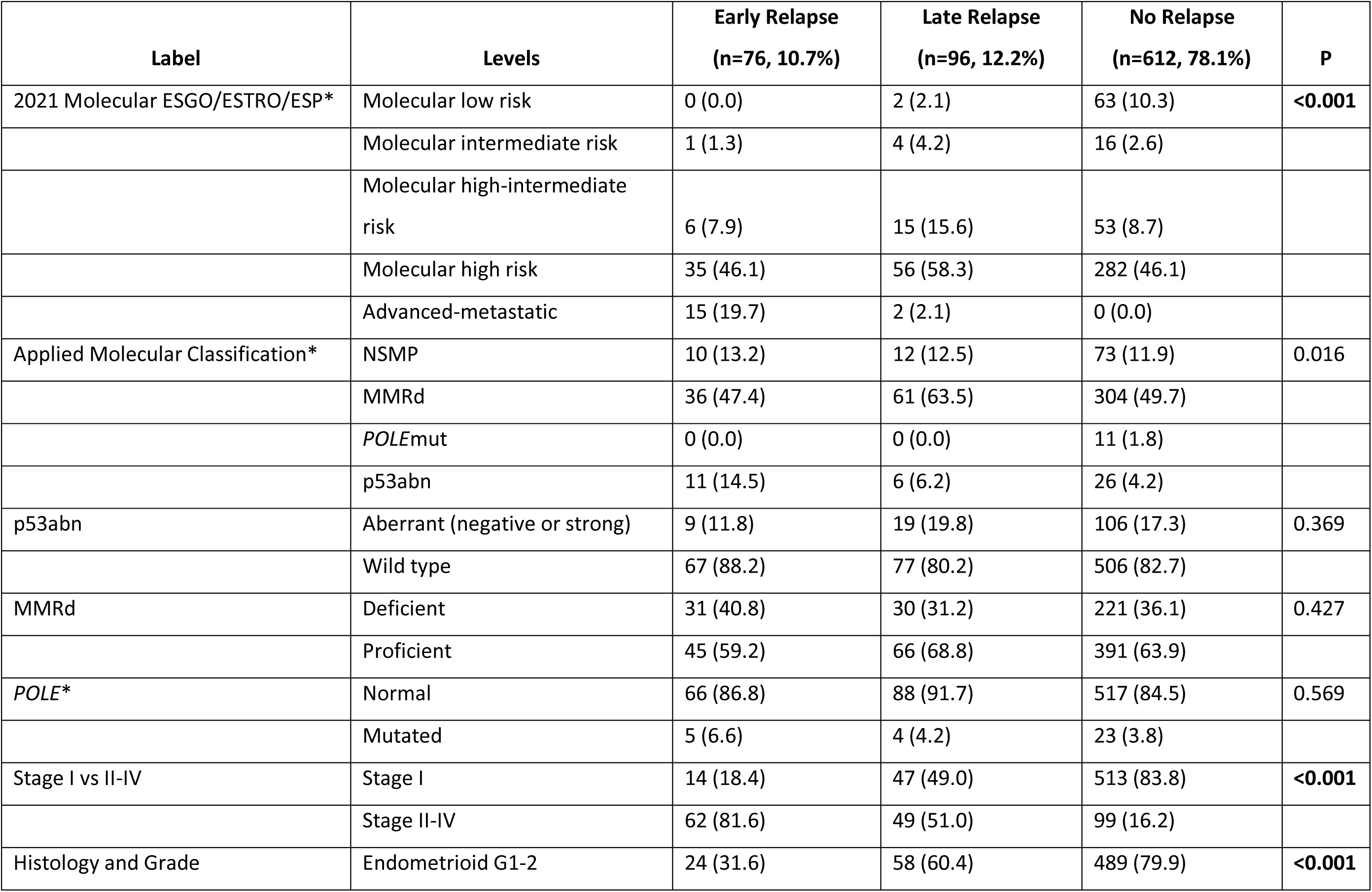
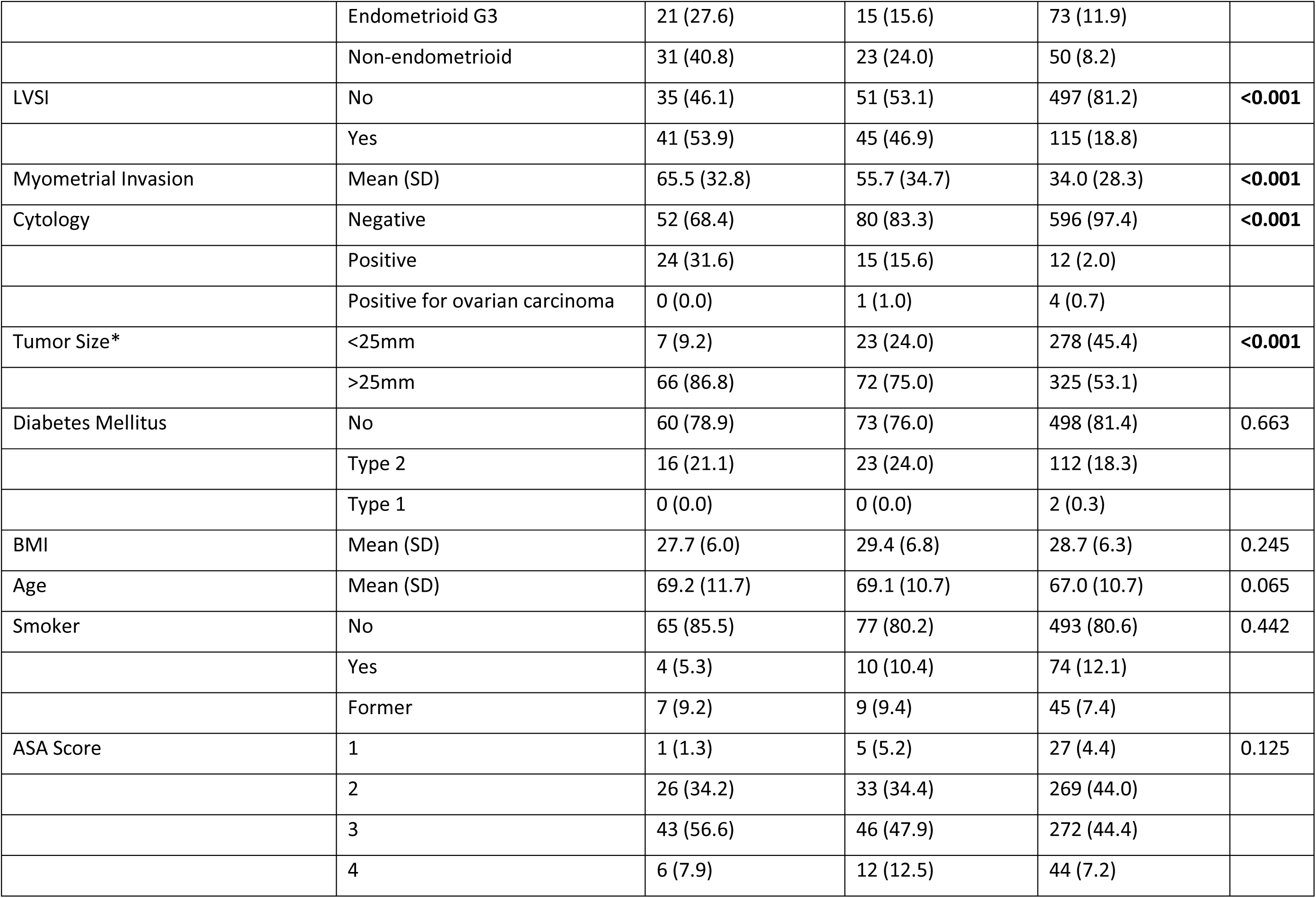
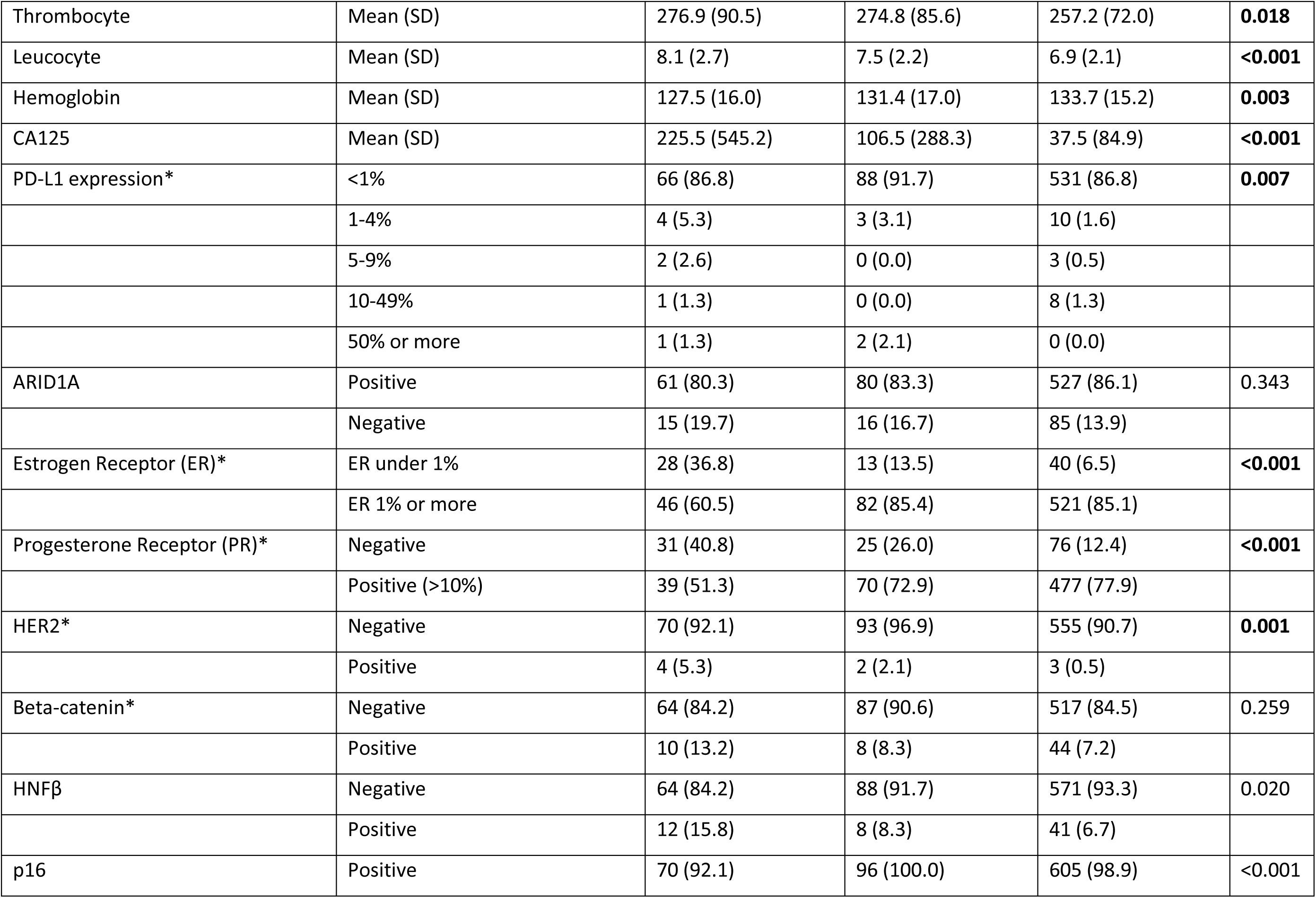
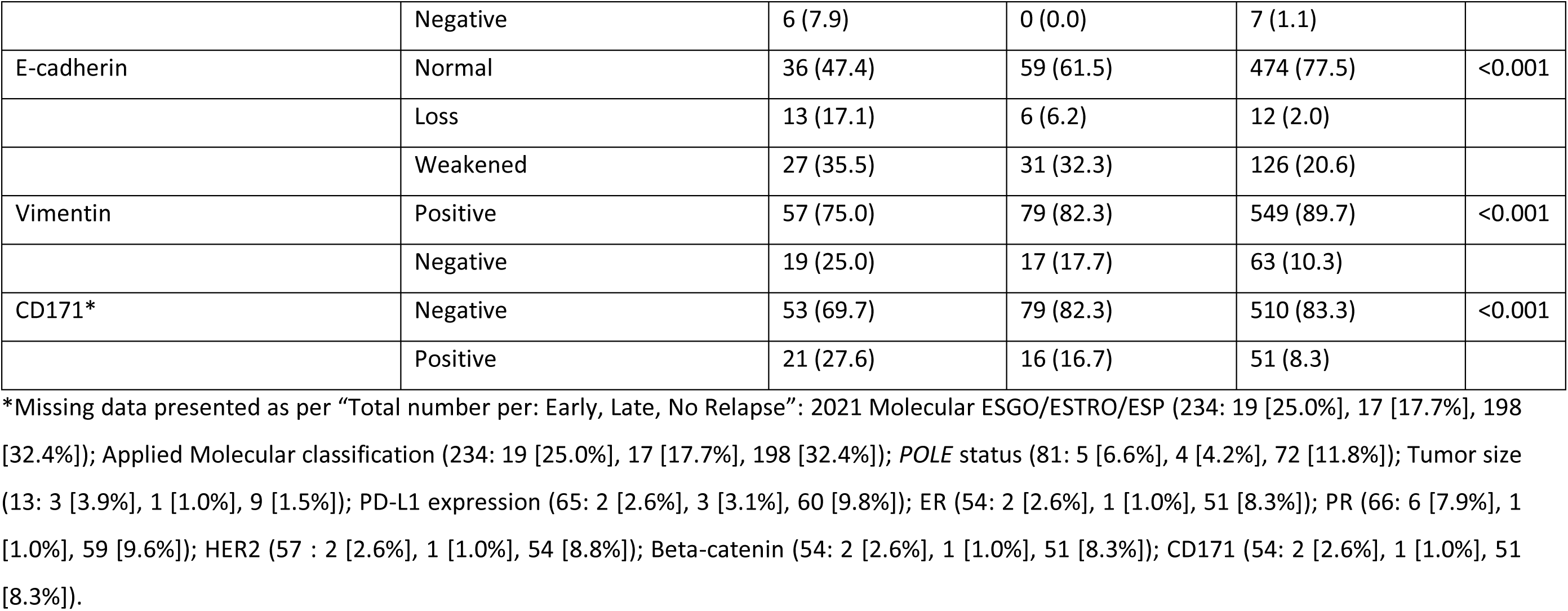
Demographic and clinicopathological characteristics of patients (n=784).

### 3.2 Feature Selection and Model Inputs

Recursive Feature Elimination (RFE) identified the key preoperative features for each model, with systemic biomarkers (CA125, thrombocytes, leucocytes) and invasion-related variables (myometrial depth, LVSI, cytology) consistently ranked highest. Traditional risk modifiers (BMI, diabetes, smoking) were rarely selected. For the risk of relapse, the TP53 + MMRd model achieved a theoretical F1 score of 0.512 using 19 features, including myometrial invasion, CA125, advanced stage (II–IV), thrombocyte count, and histology. Similar performance was observed with the applied molecular classification (F1 score = 0.492, 22 features) and the ESGO molecular classification (F1 score = 0.528, 23 features), where overlapping predictors included CA125, myometrial invasion, stage, and leucocyte count. The *POLE*-specific model achieved the highest F1 score among single-subtype analyses (0.557 with 28 features), although feature selection was limited by the sample size of *POLE*mut tumours. **Supplementary Table S2** provides detailed information on the F1 score, the number of features, and their respective names for each molecular classification.

In summary, across all analyses, systemic biomarkers (CA125, thrombocytes, leucocytes) and invasion-related variables (myometrial depth, LVSI, cytology) consistently emerged among the top-ranked features, underscoring their value for preoperative risk stratification. In contrast, traditional risk modifiers such as BMI, diabetes, and smoking status were rarely selected, suggesting their limited predictive contribution in molecularly stratified cohorts. Following feature selection, the optimal subsets of covariates retained to explain relapse risk were as follows: 19 for the TP53 + MMRd model, 22 for the molecular classification model, 28 for the POLE model, and 23 for the ESGO model (**Supplementary Table S2**).

### 3.3 Model Metrics Across the Predictive Model

### 3.4 Model Performance Comparison

Four ML models (RF, SVM, KNN, GBM) were trained across the four stratification strategies (Traditional, ESGO, TP53 + MMRd, and POLE). Performance metrics for the above-mentioned ML models are summarized in **Table 3** and visualized in **Figure 1A**. The POLE model demonstrated the highest discrimination (AUC 0.842) and sensitivity (0.886). In contrast, the Traditional model achieved the best overall accuracy (0.797) and tied with POLE for the top F1 score (0.892), while requiring fewer predictors (22 vs. 28). The TP53 + MMRd model utilized the fewest predictors (19) and delivered competitive sensitivity (0.876) and F1 score (0.872), albeit with slightly lower accuracy (0.759) and AUC (0.804). By contrast, the ESGO model underperformed, with lower discrimination (AUC 0.624) and F1 score (0.519), despite a comparable feature count (23). In summary, these findings highlight a trade-off: the Traditional model delivers a parsimonious and well-balanced solution, whereas the POLE model lays emphasis on sensitivity and discrimination. The TP53 + MMRd model offers a compact, intermediate option. Future updates will incorporate PR-AUC and balanced accuracy once harmonized per-class outputs are accessible. The detailed data for all predictive models with all four ML algorithms are outlined in **Supplementary Table S3.**

**Figure 1.**
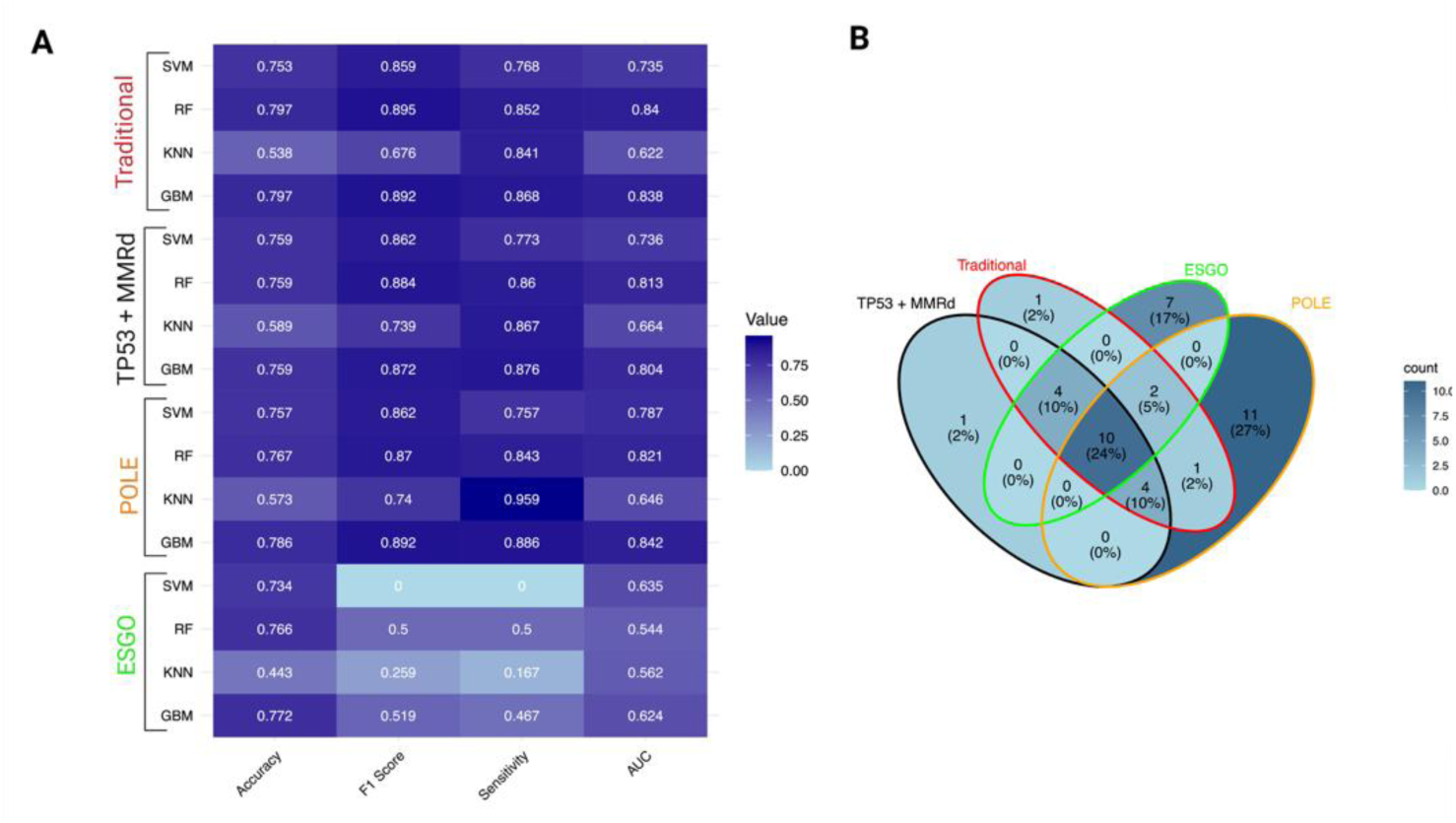
Performance Metrics and Feature Overlap Across Predictive Models. **(A)** Heatmap showing performance metrics (Accuracy, F1 Score, Sensitivity, AUC) for four machine learning algorithms (SVM, RF, KNN, GBM) across molecular subgroups: Traditional (red), TP53 + MMRd (black), POLE (orange), and ESGO (green). Darker shades indicate higher metric values. (**B)** Venn diagram illustrates the distribution of overlapping features among the four molecular subgroups. Each segment is annotated with case counts and their percentage representation. A blue gradient indicates density, highlighting both unique and shared cases across subgroup intersections.

**Table 3:**
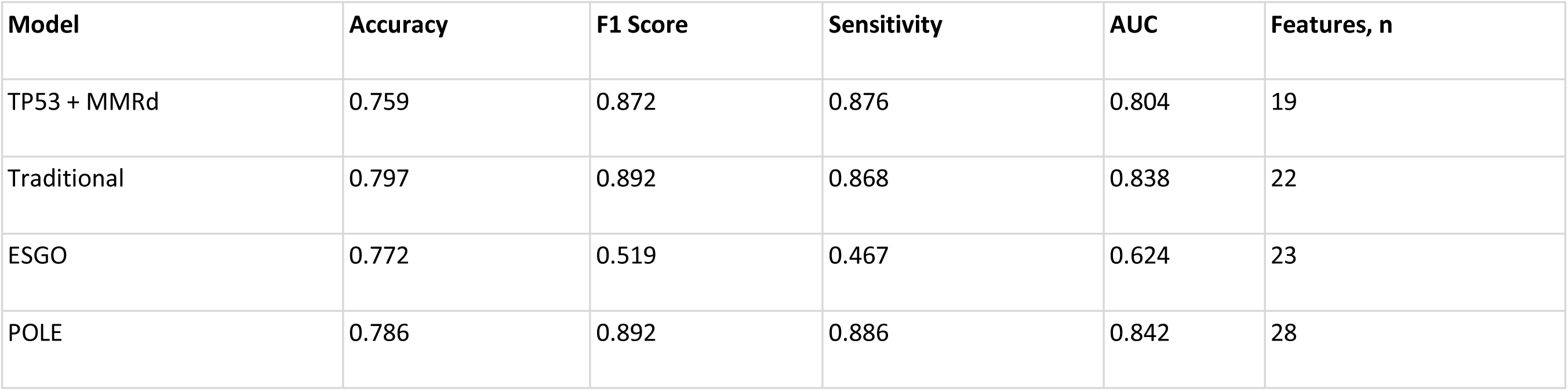
Performance of pre-operative predictive models optimized with the best Gradient Boosting (GBM) algorithm.

### 3.5 Overlap and Feature Distribution Across Models

Venn diagram analysis revealed overlapping and unique case distributions across models (**Figure 1B**). The POLE group had the highest number of cases (11), characterized by features such as LVSI, large tumor size, higher ASA score and BMI, variable PD-L1 expression (1–>10%), diabetes mellitus, and positivity for β-catenin, HNFβ, and vimentin. The ESGO group included 7 cases, mainly showing advanced or metastatic status, thrombocytosis, molecular high or intermediate risk, ER positivity, loss of E-cadherin, and a history of smoking. The TP53 + MMRd and Traditional groups each had 1 case, associated with proficient MMRd and p53abn, respectively.

Among overlapping categories, Traditional/POLE (1 case) showed positive ARID1A expression. The TP53 + MMRd/Traditional/ESGO group (4 cases) demonstrated LVSI, higher BMI and ASA scores, and larger tumor size. Similarly, TP53 + MMRd/Traditional/POLE (4 cases) was characterized by ER positivity, thrombocytosis, normal E-cadherin, and non-smoking status. The Traditional/ESGO/POLE overlap (2 cases) involved endometrioid G3 tumors with weakened E-cadherin. The most complex intersection, TP53 + MMRd/Traditional/ESGO/POLE (10 cases), showed high-risk clinicopathologic features including stage II–IV disease, myometrial invasion, elevated CA125, positive CD171 and cytology, older age, leukocytosis, PR positivity, low hemoglobin, and non-endometrioid histology. Full details about the feature are presented in **Supplementary Table S4.**

### 3.6 Traditional Model: Class-Specific Performance

Per-class performance metrics for the traditional models revealed the expected imbalance pattern (**Figure 2A)**. The model demonstrated optimal performance for the No-Relapse class (F1=0.892, precision=0.868, recall=0.918, PR-AUC=0.935; accuracy=0.829). Performance for the Early Relapse class was moderate (F1=0.615, precision=0.571, recall=0.667, PR-AUC=0.477; accuracy=0.937), reflecting reasonable detection with some false positives. The Late-Relapse class posed the greatest challenge (F1=0.308, precision=0.400, recall=0.250, PR-AUC=0.266; accuracy=0.829), consistent with class rarity and overlap with other phenotypes. Overall, these results support the importance of reporting PR-AUC and F1 alongside ROC-AUC, and they suggest the implementation of targeted strategies, such as rebalancing or multimodal features, to enhance minority-class detection. Per-class performance of the Traditional model across relapse timing is presented in **Supplementary Table S5.**

**Figure 2.**
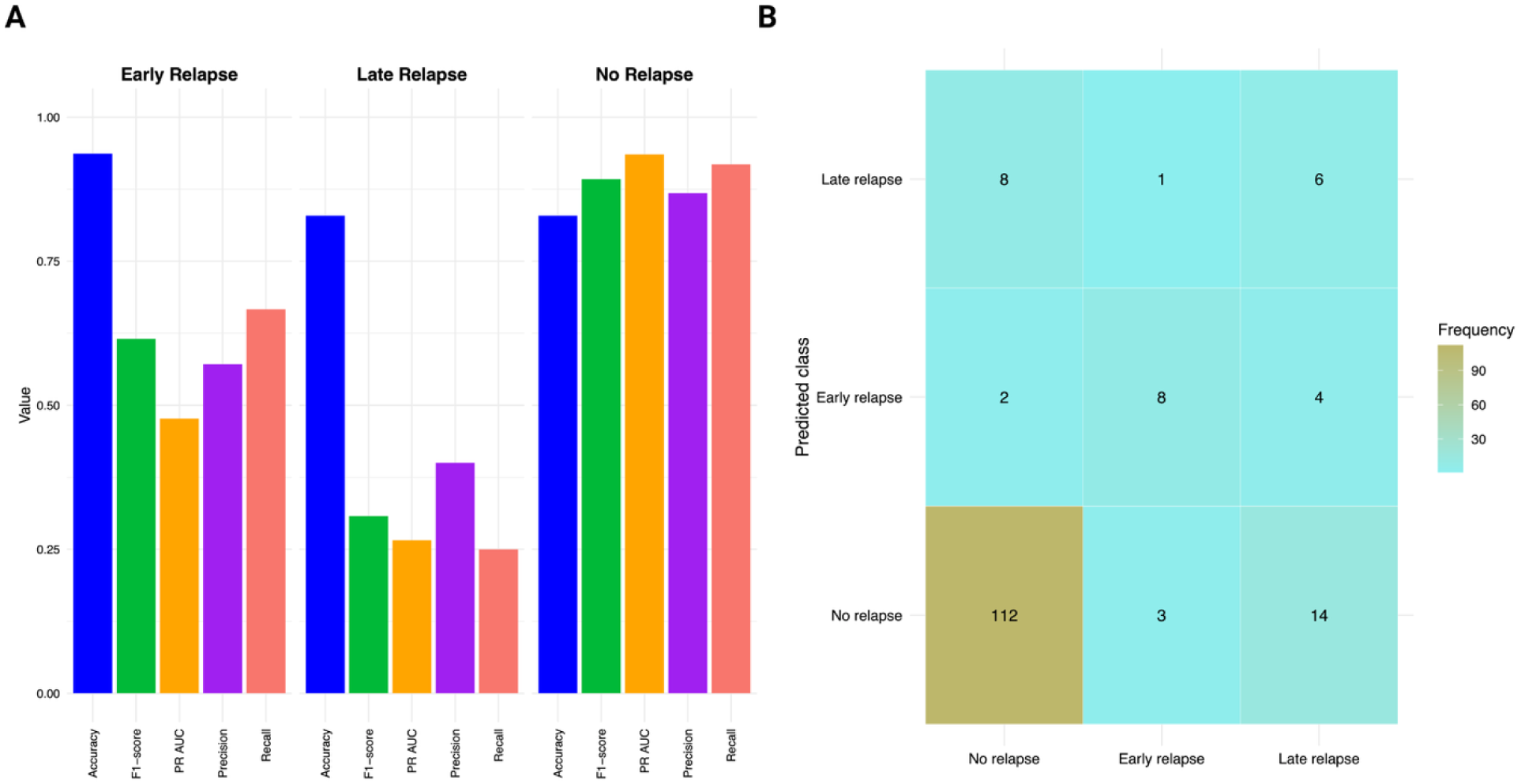
Class-Level Performance and Confusion Matrix for the Traditional Model. **(A)** Grouped bars display accuracy, F1, PR-AUC, precision, and recall for Early Relapse, Late Relapse, and No Relapse classes. The highest performance is observed for the No-Relapse class (F1 0.892; PR-AUC 0.935), intermediate for Early Relapse (F1 0.615; PR-AUC 0.477), and lowest for Late Relapse (F1 0.308; PR-AUC 0.266). These patterns reflect both the underlying class imbalance and the greater challenge of distinguishing Late Relapse events. **(B)** Confusion matrix for relapse stages (No-Relapse, Early Relapse, Late Relapse) with row-normalized recall percentages. Rows indicate true labels, while columns indicate predicted labels. The recall is strong for No-Relapse, moderate for Early Relapse (often misclassified as No-Relapse), and weak for Late Relapse (mostly misclassified as No-Relapse). Darker colors indicate a higher frequency of recall.

The confusion matrix corroborates these class-specific metrics (**Figure 2B**). Notably, the’No-Relapse’ category predominantly occupies the diagonal, with most instances correctly classified. The’Early Relapse’ category exhibits a moderate true-positive block, with most remaining errors misclassified as’No-Relapse.’ The’Late Relapse’ category has the least populated true-positive cell, with misclassifications primarily occurring as’No-Relapse’ and, to a lesser extent, as’Early Relapse.’ This asymmetric error pattern (Late → Early/No) aligns with class imbalance and the lower recall for late events, whereas early events are detected with greater reliability, and’No-Relapse’ is identified with high recall.

### 3.7 XAI-Based Model Interpretability: Traditional Model

SHAP analyses demonstrated class-specific patterns that aligned with the overall feature ranking. The global importance was highlighted by the mean absolute SHAP values (organized by class), which pinpointed FIGO stage, E-cadherin status, tumor size, and LVSI as the primary predictors. These were followed by ARID1A, PR, and peritoneal cytology. There were also smaller contributions from hematologic/host factors such as thrombocytes, BMI, leucocytes, and hemoglobin, as well as CA125, myometrial invasion, and ASA score. Notably, these variables had varying levels of influence across different classes, as illustrated by the stacked bar profiles **(Figure 3A).**

**Figure 3.**
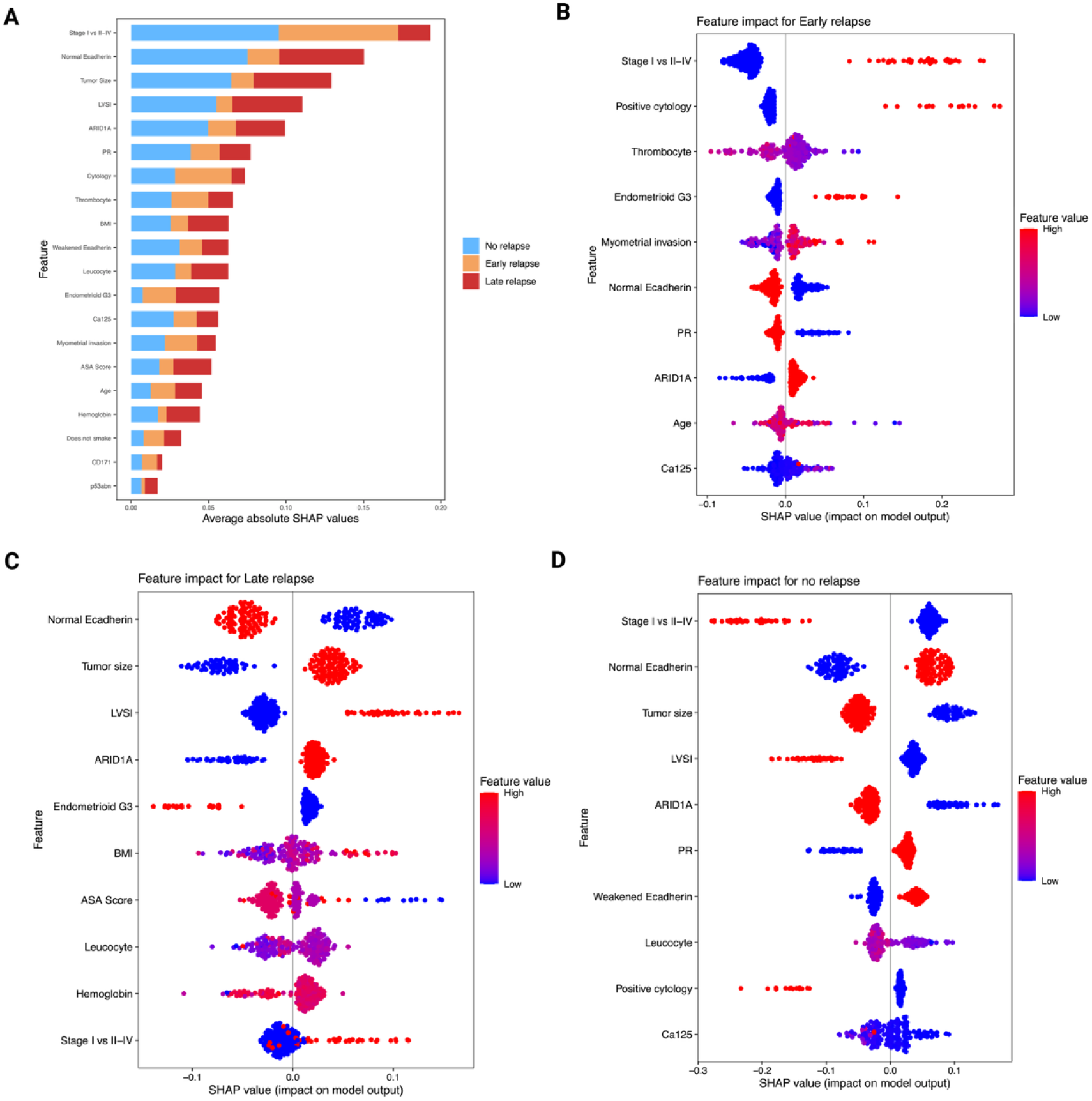
SHAP-Based Interpretation of Relapse Risk in the Traditional Model. **(A)** Global SHAP summary plot showing mean absolute SHAP values, ranked by feature importance and stacked by relapse class. This visualization highlights the most influential predictors across all classes. **(B–D)** Class-specific SHAP beeswarm plots for Early Relapse, Late Relapse, and No Relapse, respectively. Each point represents an individual patient; color indicates the feature value (red = high, blue = low), while horizontal position reflects the magnitude and direction of the feature’s impact on the predicted class probability. These plots illustrate how specific features contribute to relapse risk stratification at the patient level.

Early Relapse (≤6 months) was primarily driven by higher stage (II–IV), positive peritoneal cytology, thrombocytosis, grade-3 endometrioid histology (G3), deep myometrial invasion, and ARID1A loss, with elevated CA125 providing additional risk. Protective features included PR positivity (>10%), preserved E-cadherin, and smaller tumor size **(Figure 3B).** Late-relapse class (>6 months) was most strongly linked to LVSI positivity, larger tumor size, and higher stage, supported by ARID1A loss, G3 endometrioid histology, and host factors such as higher BMI/ASA, leukocytosis, and lower hemoglobin. In the Late-Relapse class, Stage I status, preserved E-cadherin, and smaller tumor size served as protective features, each contributing negative SHAP values. The latter implies that they decreased the model’s log-odds and predicted probability of Late Relapse (shifting probability toward Early/No Relapse) when other inputs were held constant **(Figure 3C)**. In the No-Relapse category, factors that favored predictions of No Relapse included Stage I disease, preserved E-cadherin, PR levels above 10%, low CA125, negative cytology, absence of LVSI, and smaller tumors. In contrast, Stage II–IV, positive LVSI, ARID1A loss, larger tumors, and positive cytology decreased the likelihood of remaining free from recurrence. Early Relapse is associated with tumor burden and biological aggressiveness, such as advanced stage, positive cytology, elevated CA125, and invasion/size. Late Relapse is more closely linked to anatomic spread and size, including advanced LVSI, larger diameter, and higher stage, while No-Relapse reflects the opposite profile **(Figure 3D)**. Collectively, the panels provide a coherent and clinically intuitive distinction of relapse phenotypes. Early Relapse is primarily driven by tumour biology and burden, while Late Relapse is more associated with anatomic spread and size. In contrast, the No-Relapse profile reflects the inverse of these features. This alignment between SHAP-based interpretability and established clinical expectations offers clinical plausibility and bolsters confidence in the model’s predictive framework.

### 3.8 POLE Model: Class-Specific Performance

Per-class performance metrics for the POLE model revealed a performance pattern consistent with the traditional model, largely influenced by class imbalance **(Figure 4A).** The model achieved its strongest performance for the No-Relapse class (F1 = 0.892, Precision = 0.886, recall = 0.897, PR-AUC = 0.944; accuracy = 0.835), indicating stable generalization and consistent identification of patients without relapse. The Early Relapse class showed moderate discrimination (F1 = 0.593, Precision = 0.533, recall = 0.667, PR-AUC = 0.649; accuracy = 0.893), indicating a balanced sensitivity, precision, with a modest improvement in PR-AUC, compared to the traditional model. The Late Relapse class exhibited the lower performance (F1 = 0.273, Precision = 0.333, recall = 0.231, PR-AUC = 0.208; accuracy = 0.845), reflecting continued limitations in minority-class detection. Collectively, these findings reinforce the need for targeted modeling strategies such as temporal reweighting or synthetic data augmentation to enhance recognition of rare relapse patterns, particularly late events. Per-class performance of the POLE model across relapse timing is presented in **Supplemental Table S5.**

**Figure 4.**
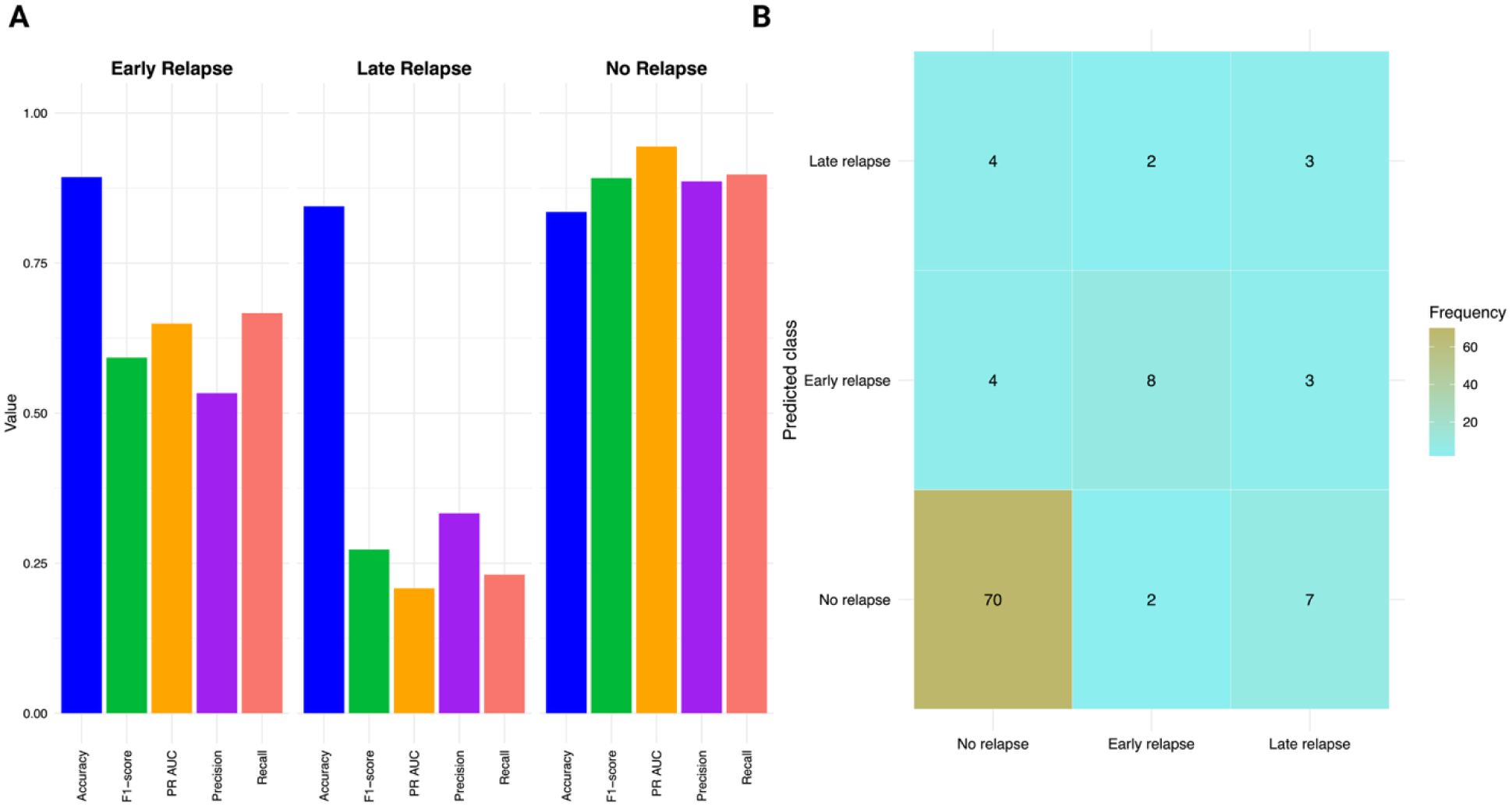
Class-Level Performance and Confusion Matrix for the POLE Model. **(A)** Grouped bar chart displaying performance metrics — Accuracy, F1 Score, Precision-Recall AUC (PR-AUC), Precision, and Recall for each relapse class: Early Relapse, Late Relapse, and No Relapse. The model showed highest performance for the No-Relapse class (F1 = 0.891; PR-AUC = 0.944), moderate performance for Early Relapse (F1 = 0.592; PR-AUC = 0.649), and lowest for Late Relapse (F1 = 0.272; PR-AUC = 0.208). These results reflect both class imbalance and the increased difficulty in identifying Late Relapse events. (**B)** Confusion matrix for the POLE model predictions (Early Relapse, Late Relapse, No Relapse), displaying row-normalized recall percentages. Rows correspond to true labels and columns to predicted labels. Recall is strongest for No Relapse, moderate for Early Relapse (often misclassified as No Relapse), and lowest for Late Relapse, which are commonly misclassified as Early or No Relapse. Darker shading indicates higher recall frequency.

The confusion matrix for the POLE model (**Figure 4B**) illustrates the class-specific trends. Notably, the recall was strongest for the No-Relapse class, moderate for Early Relapse (often misclassified as No Relapse), and weakest for Late Relapse (frequently confused with both Early and No Relapse). The dominance of correct No-Relapse predictions forms a dense diagonal cluster, while the asymmetric misclassification of Late Relapse underscores both data imbalance and model conservatism. Despite this, the slightly higher PR-AUC for Early Relapse suggests improved precision–recall trade-offs relative to the traditional model.

### 3.9 XAI-Based Model Interpretability: POLE Model

The SHAP patterns for the POLE models (**Figure 5**) closely resemble those of the Traditional model (**Figure 3**). On a global scale, FIGO stage, tumor size, LVSI, and myometrial invasion emerged as the primary contributors by mean SHAP **(Figure 5A).** Conversely, CA125 and peritoneal cytology were associated with increased risk, while progesterone receptor (PR) >10% shifted predictions towards No Relapse **(Figure 5B).** In terms of class-specific SHAP, Early Relapse was associated with higher stage, positive cytology, and elevated CA125 levels. In contrast, Late Relapse was linked to positive LVSI and larger tumor size **(Figure 5C).** Tumor stage I preserved E-cadherin, and smaller tumor sizes acted as protective factors against Late Relapse, showing negative SHAP contributions that reduced the model’s probability of Late Relapse and directed predictions toward No Relapse **(Figure 5D).** Compared to the Traditional model, POLE exhibited a slight reweighting, highlighting the influence of size/LVSI for Late Relapse and cytology/CA125 for Early Relapse, while maintaining the early versus late pattern.

**Figure 5.**
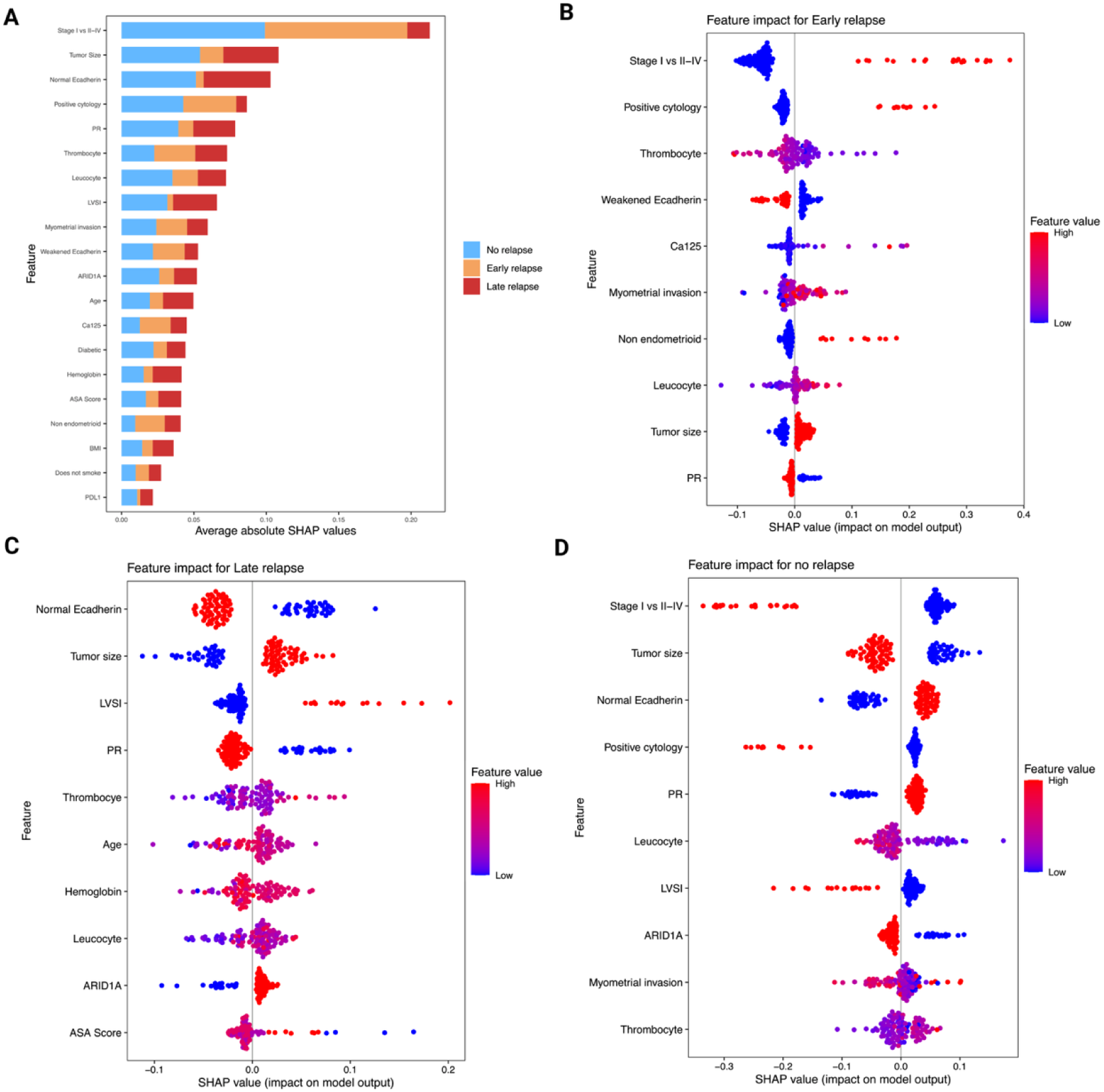
SHAP-Based Interpretation of Relapse Risk in the POLE Model. **(A)** Global SHAP summary plot displaying mean absolute SHAP values, ranked by overall feature importance and stacked by relapse class. This visualization highlights the most influential predictors contributing to relapse risk across the POLE subgroup. **(B–D)** Class-specific SHAP beeswarm plots for Early Relapse, Late Relapse, and No Relapse, respectively. Each point represents an individual patient; color indicates the feature value (red = high, blue = low), while horizontal position reflects the magnitude and direction of the feature’s impact on the predicted class probability. These plots provide insight into how specific features drive model predictions at the patient level, supporting individualized risk assessment.

## 4. Discussion

### 4.1 Study Overview and Clinical Context

This study demonstrates that preoperative multimodal data, when analyzed through interpretable ML, can effectively predict relapse risk and timing in EC. By analyzing a large, molecularly stratified cohort of 784 patients, we developed four complementary ML models that integrate clinicopathological, molecular, and systemic features. These models offer actionable insights for individualized risk stratification prior to surgery or definitive histopathological staging.

### 4.2 Model Performance and Biological Interpretability

Among all the molecular models, the Traditional model achieved the highest overall accuracy (0.797) and demonstrated balanced performance, while the POLE-based model excelled in sensitivity (0.886) and discrimination (AUC 0.842). The TP53 + MMRd model, which focuses on high-risk molecular subsets, maintained competitive performance with fewer variables, underscoring the biological significance of molecularly driven compact modeling. In contrast, the ESGO-based classifier exhibited limited discrimination (AUC 0.624), indicating that static risk categories may underrepresent the complexity of relapse heterogeneity.

Our SHAP analyses revealed that relapse timing reflects distinct biological patterns: Early Relapse is primarily driven by tumor burden and systemic inflammation (advanced stage, positive cytology, high CA125, thrombocytosis), whereas Late Relapse is associated with invasion-related features (LVSI, tumor size, ARID1A loss). Predictions of No Relapse were characterized by PR receptor positivity, preserved E-cadherin, and early-stage disease, reinforcing the biological coherence of the model outputs. This mechanistic interpretability enhances clinical confidence and provides actionable decision support.

### 4.3 Comparison with Existing Literature

Prior prognostic tools in EC have mainly relied on postoperative clinicopathological variables such as FIGO stage, tumor grade, and LVSI, which inadequately capture the heterogeneity of high-risk molecular subtypes (Concin et al., 2021; Oaknin et al., 2022). Incorporation of TCGA-based Molecular classifiers into WHO/ESGO/FIGO systems has markedly improved risk stratification, yet predictive accuracy for relapse remains suboptimal, particularly in NSMP, p53abn, and carcinosarcoma cases (León-Castillo et al., 2025).

In fact, molecular context significantly influences prognosis. The ProMisE classifiers enable the practical implementation of TCGA subgroups (Talhouk et al., 2015), and PORTEC-3 showed that p53abn cases, while associated with poor outcomes, benefited from chemoradiotherapy over radiotherapy alone (de Boer et al., 2019). Recent cohort studies further confirmed distinct relapse patterns across molecular subtypes, reinforcing the need for molecularly informed prediction models (Lindemann et al., 2025; Loukovaara et al., 2025). Notably, semiquantitative LVSI remains prognostically relevant within endometrioid EC regardless of molecular classification (Loukovaara et al., 2025b), suggesting that traditional histopathological markers still hold relevance when interpreted in a molecular context, consistent with our findings. Interestingly, the overlap and feature distribution across models highlight both shared and distinct prognostic profiles. Notably, the POLE group, despite its generally favorable prognosis (Liu et al., 2017), exhibited several high-risk features such as LVSI, elevated BMI and ASA scores, and variable PD-L1 expressions, suggesting that even molecularly favorable subtypes may harbor complex clinicopathologic traits. The ESGO group, enriched for advanced disease and molecular high-risk status, aligns with prior findings linking ER positivity and E-cadherin loss to aggressive behavior (Murali et al., 2019). Overlapping categories, particularly the TP53 + MMRd/Traditional/ESGO/POLE cluster, revealed a convergence of high-risk features, including stage II–IV disease, myometrial invasion, and elevated CA125 — traits previously associated with poor outcomes (de Boer et al., 2019). These intersections underscore the value of multimodal profiling in capturing relapse risk beyond single-model stratification.

Recent ML models such as TJHPEC (Wang et al., 2022), NU-CATS (Zheng et al., 2023), and im4MEC (Fremond et al., 2023) have shown promise but often lack integration of multimodal biomarkers or preoperative applicability. While HECTOR integrated whole-slide histopathology with stage across eight EC cohorts (including PORTEC trials) to deliver strong prognostic performance with therapy-relevant stratification (Volinsky-Fremond et al., 2024), our framework complements these efforts by offering infrastructure-light, interpretable predictions at the patient level. Similarly, a large multi-institutional Israeli XGBoost model (n≈1,935) demonstrated feasibility with SHAP-informed relapse prediction (AUC ≈ 0.84) (Ohad Houri et al., 2022), while MRI radiomics models integrating intertumoral and peritumoral features predict relapse with decision-curve utility (Li et al., 2025). Beyond EC, ML-based relapse prediction in breast cancer has shown that hybrid mechanistic and ML models improve calibration for Late Relapses (Nicolò et al., 2020), suggesting opportunities for similar approaches in EC. To our knowledge, by incorporating systemic and immunohistochemical markers alongside molecular classification, our approach aligns with emerging evidence supporting the value of multimodal inputs for relapse prediction (Karpel et al., 2023; Njoku et al., 2022). Notably, the identification of ARID1A and p16 as key relapse predictors aligns with prior studies linking these biomarkers to aggressive EC phenotypes (Liu et al., 2017; Murali et al., 2019).

### 4.4 Interpretability and Clinical Implications

SHAP-based interpretability provided transparent, class-specific insights into model predictions by highlighting key biomarkers and risk factors, such as ARID1A loss, elevated CA125, thrombocytosis, p16 expression, FIGO stage, LVSI, cytology, tumor size, and E-cadherin status. These features aligned well with established molecular and staging frameworks, reinforcing the biological plausibility of the models.

Importantly, SHAP profiles enabled differentiation between relapse phenotypes. Patients flagged as high risk for Early Relapses, often driven by tumor burden and systemic inflammation, may benefit from intensified imaging or systemic therapy. Conversely, those with Late-Relapse signals, typically associated with anatomic spread, may require extended surveillance. This level of interpretability supports personalized treatment planning and enhances clinical trust in ML-based decision support.

Embedding these models into electronic health records and generating patient-level risk summaries could facilitate their integration into multidisciplinary care. This approach is consistent with evolving ESGO– ESTRO–ESP and FIGO 2023 guidelines, which increasingly emphasize molecular stratification in EC management (Berek et al., 2023; Concin et al., 2025; Maria-Bianca Anca-Stanciu et al., 2025).

For aggressive subtypes such as carcinosarcoma, which consistently showed high predicted relapse risk and distinct biomarker profiles, these models may be particularly impactful. Their clinical utility could mirror precision strategies already in use, such as trastuzumab for HER2-positive uterine serous carcinoma (Fader et al., 2020; Fader et al., 2018), demonstrating how biomarker-informed ML tools can personalize care and improve outcomes.

Our models achieved an AUC of up to 0.842, comparable to the externally validated ENDORISK-2 framework for predicting lymph-node metastasis in EC (AUC ≈ 0.85). Both approaches highlight the value of integrating molecular and clinicopathological data in preoperative assessment. While ENDORISK-2 informs nodal management, our models focus on relapse timing, offering complementary guidance for early treatment planning and surveillance before surgery (Lombaers et al., 2025).

### 4.5 Strengths and Limitations

The key strengths of this study include the use of a large, well-characterized cohort, exclusive reliance on preoperative data, integration of systemic, immunohistochemical, and molecular data, and explainable AI outputs ensuring clinical interpretability. Limitations include the single-center retrospective design and class imbalance affecting Late Relapse prediction. Additionally, while SHAP improves interpretability, it is sensitive to feature selection and model architecture, which may influence the stability and generalizability of feature importance rankings (Ponce-Bobadilla et al., 2024). External validation and prospective integration into clinical workflows are needed to confirm generalizability.

### 4.6 Conclusions and Future Directions

We developed preoperative, interpretable ML models capable of predicting relapse timing in EC, aligning with molecular risk frameworks. Stage, LVSI, tumor size, myometrial invasion, CA125, cytology, and hormone-receptor status emerged as key predictors across algorithms, showing clinically coherent effects. Gradient boosting with SHAP explanations provides patient-level transparency, supporting decision-making for surveillance and adjuvant therapy, particularly in molecularly adverse groups and aggressive histology.

To build on these findings, future work should focus on validating the models in external, multi-center cohorts and integrating them into prospective clinical workflow. Combining clinicopathological data with radiomics, liquid biopsy, and whole-slide pathology could enhance predictive accuracy (e.g., HECTOR-like and radiomics pipelines) and enable trial-based validation (Li et al., 2025; Volinsky-Fremond et al., 2024). Additionally, incorporating relapse site prediction, such as regional versus distant recurrence, could further refine surveillance strategies and therapeutic planning. Adaptive learning frameworks may allow real-time updates as new data becomes available, improving clinical responsiveness. Embedding explainable ML tools into electronic health records and developing clinician-friendly interfaces will be key to adoption. Finally, ethical considerations such as transparency, patient communication, and equitable access must be addressed to ensure responsible implementation in oncological care.

## Credit Authorship Contribution Statement

Conceptualization, investigation, methodology: SVM, MK, AP, RB, AS, ML, VM. Data curation, Resources: MK, ML, AP, RB. Formal analysis: SVM, VM. Visualization, investigation: SVM, MK, VM. Writing – original draft preparation: SVM, MK, VM. Writing – review & editing: SVM, MK, AP, RB, AS, ML, VM. Supervision: VM, ML. Funding acquisition, project administration: RB, AS, VM.

## Code Availability

The scripts used to perform the different analyses described in this paper can be found in https://github.com/SergioVela17/Preoperative-Relapse-Prediction-in-Molecularly-Stratified-Endometrial-Cancer-Finnish-Cohort-Study.

## Acknowledgement

We thank Annikki Löfhjelm for the technical support.

## Ethics Statement

Ethical approval was obtained from the Helsinki University Hospital Institutional Review Board (HUS/491/2021) and the Finnish Medicines Agency (FIMEA/2021/005153).

## Consent for Publication

Informed consent was waived for this retrospective cohort.

## Supplementary Files

Supplementary Table S1. Details of immunohistochemistry, scoring, and interpretation Supplementary Table S2. Distribution of Recursive feature elimination (RFE) derived data

Supplementary Table S3. Performance metrix of pre-operative predictive models derived from all ML algorithm.

Supplementary Table S4. Overlapping molecular and clinical features across the predictive model via Venn diagram.

Supplementary Table S5. Per-class performance metrics for the Traditional and POLE model

## Data Availability

The datasets used and/or analyzed during the current study are available from the corresponding author on reasonable request.

